# A deep learning pipeline for segmentation of *Proteus mirabilis* colony patterns

**DOI:** 10.1101/2022.01.17.475672

**Authors:** Anjali Doshi, Marian Shaw, Ruxandra Tonea, Rosalía Minyety, Soonhee Moon, Andrew Laine, Jia Guo, Tal Danino

## Abstract

The motility mechanisms of microorganisms are critical virulence factors, enabling their spread and survival during infection. Motility is frequently characterized by qualitative analysis of macroscopic colonies, yet the standard quantification method has mainly been limited to manual measurement. Recent studies have applied deep learning for classification and segmentation of specific microbial species in microscopic images, but less work has focused on macroscopic colony analysis. Here, we advance computational tools for analyzing colonies of *Proteus mirabilis*, a bacterium that produces a macroscopic bullseye-like pattern via periodic swarming, a process implicated in its virulence. We present a dual-task pipeline for segmenting (1) the macroscopic colony including faint outer swarm rings, and (2) internal ring boundaries, unique features of oscillatory swarming. Our convolutional neural network for patch-based colony segmentation and U-Net with a VGG-11 encoder for ring boundary segmentation achieved test Dice scores of 93.28% and 83.24%, respectively. The predicted masks at times improved on the ground truths from our automated annotation algorithms. We demonstrate how application of our pipeline to a typical swarming assay enables ease of colony analysis and precise measurements of more complex pattern features than those which have been historically quantified.

## 1. INTRODUCTION

Bacteria colony development processes, such as swarming motility, have been implicated in the pathogenicity of many microorganisms, enabling their spread and survival in unfavorable conditions, such as in the presence of antimicrobials [1-4]. Motility is studied not only on the microscopic scale, but also the macroscopic scale via colony growth assays under conditions that produce different types of motility. Analysis is traditionally laborious, time-consuming, and low-throughput, often involving qualitative comparison or manual measurement of individual colonies [5, 6]. Recent advancements in image acquisition, image processing, and computer vision can enable easier, scalable, and nuanced analysis of colony features. However, these advances have mainly been applied to microscopic images of a limited set of microbial species [7-9]. A few recent studies have begun the work on the macroscopic scale, but they have analyzed colonies with well-defined contours and relatively simple inner features [10, 11]. Many relevant species have more complex colonies with unique internal features and poorly-defined boundaries, generated by a variety of motility mechanisms. For example, the common soil bacterium and pathogen *Proteus mirabilis* rapidly migrates across solid surfaces via periodic swarming: a highly coordinated movement propelled by flagella. Alternating swarming with phases of rest and division, *P. mirabilis* produces a sequential array of macroscopic rings when inoculated on laboratory agar [5, 12]. The role of swarming in *P. mirabilis* infection of the lungs, wounds, and urinary tract, especially in the presence of catheters, is an area of active research [13]. Detection and measurement of the bacterium’s periodic colony features could shed more light on its virulence. These features have yet to be quantified in detail, as typical analysis involves measurement of colony radii with a ruler or in ImageJ. Traditional edge detection methods are insufficient for segmentation, as depending on experimental conditions, the ring boundaries can be numerous, densely spaced, and/or indistinct, compounding the difficulty of quantification. Additionally, to our knowledge, no large datasets of *P. mirabilis* swarm colony images exist to enable development of automated approaches.

To robustly analyze swarm colony formation, we developed a semi-automated pipeline for segmentation of macroscopic *P. mirabilis* colonies and their ring boundaries, using a dataset generated in our laboratory **(Fig. 1)**. The workflow begins with data acquisition, image preprocessing, and automated annotation. Two parallel approaches using convolutional neural networks (CNNs) are implemented for colony and ring boundary segmentation. Our first approach uses a patch classification-based CNN and label fusion to segment the colony, including faint active swarm rings, from background agar. The second approach uses a VGG-11 U-Net to segment precise boundaries of a colony’s rings generated by completed swarming events and postprocessing to refine the predictions [14-16]. The two models provide sufficient information to efficiently quantify important motility features in collections of colony images. We demonstrate the utility of this pipeline by showing how it enables feature extraction from a standard assay investigating swarming under different conditions.

**Fig. 1.**
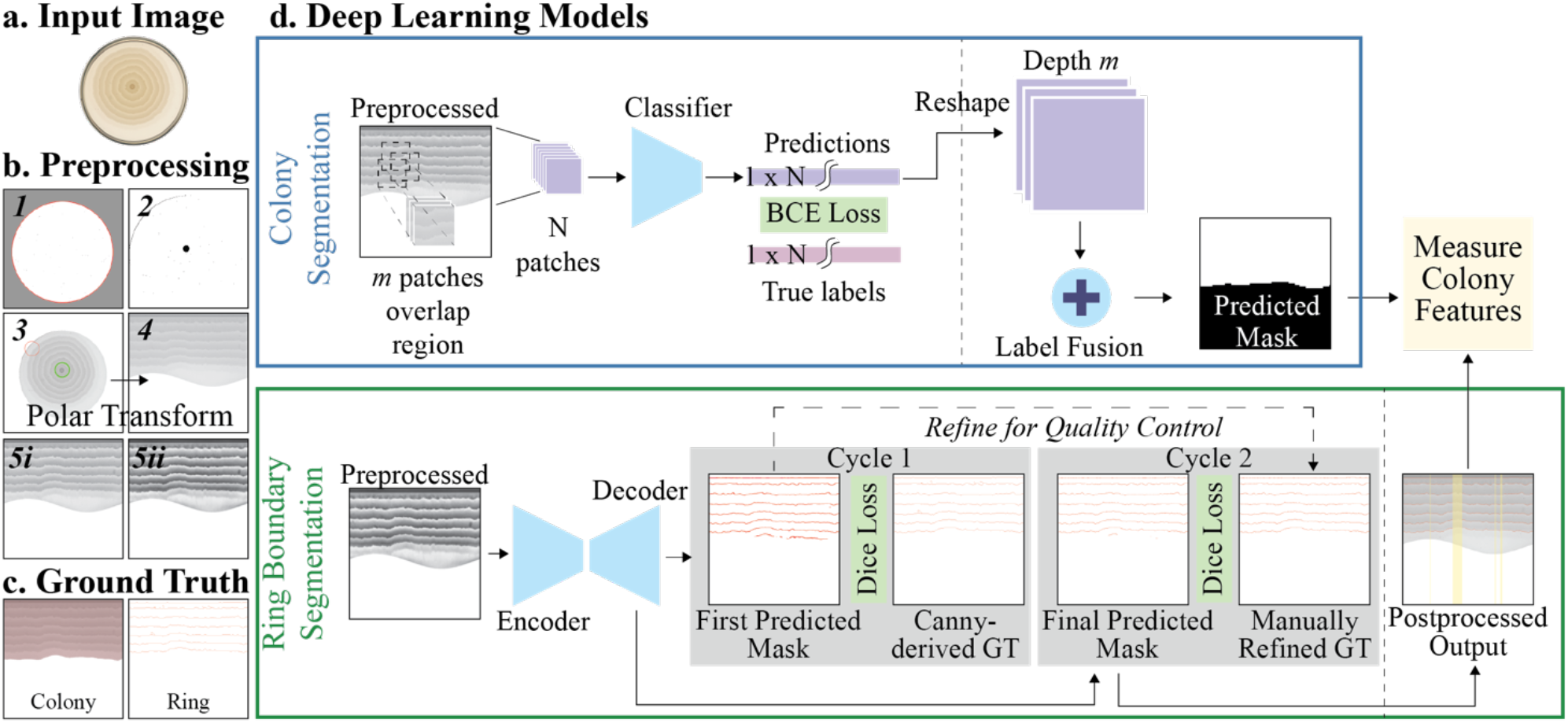
Dual-task segmentation pipeline schematic. **a-b**. RGB Cartesian coordinate images are transformed to grayscale polar coordinate form **(b1-4)**, then further preprocessed for subsequent colony **(b5i)** and ring boundary **(b5ii)** segmentation tasks. **c**. Separate algorithms generate initial ground truth approximations of colony and ring boundary masks from preprocessed images. **d**. Parallel deep learning models are trained and tested for colony and ring boundary segmentations, enabling automated feature extraction. Here, the postprocessed predicted ring boundary mask is laid over image **b4** with omitted columns (lacking the true ring boundary count) highlighted in yellow.

## 2. METHODOLOGY

We first present image acquisition and preprocessing protocols for dataset preparation **(Table 1)**. We then describe two methods for segmenting (1) bacterial colonies from background and (2) inner ring boundaries. Two overlapping datasets of images are used, chosen based on appropriateness for the given task (i.e., with both colony and agar spaces for colony segmentation, or with more distinct rings for ring segmentation). Both tasks begin with annotation procedures followed by training CNNs.

**Table 1.** Image acquisition and datasets.

| Swarm Assay | Scanner Settings | Segmentation Task | Dataset Size | Dataset Description | Model Input Type |
| --- | --- | --- | --- | --- | --- |
| 1.3-1.5% agar, 30 min plate drying, 2 uL OD 0.1 culture inoculated, 15 min drying, 24 hour growth at 25-37°C | 400 DPI, 24-bit color, Orientation: agar facing ceiling, remove plate lid | Colony | 306 | Colonies covering part or full plate, agar-only images | Image patches (150,000 positive/112,000 negative) |
|  |  | Ring Boundaries | 558 (Cycle 1), 300 (Cycle 2) | Colonies with distinct ring boundaries | Fullsize images padded (border reflect method) |

### 2.1. Swarm assays, data acquisition, and image preprocessing

Swarming assays were conducted similarly to the method presented in [17]. We maintained standard conditions throughout all assays for reproducibility, including agar volume and drying time, concentration and volume of bacteria inoculated, and the length of colony incubation **(Table 1)**. Plates were scanned using an Epson Perfection V800 Photo scanner with consistent settings, generating images of around 1500×1500 pixels. These protocols for conducting and documenting swarming assays can be easily used by different labs, requiring only readily available equipment and reagents.

Going beyond traditional manual processing of swarm images, we developed a semi-automated MATLAB script to preprocess images. After a user inputs an image directory path, each image is converted to grayscale and the Petri rim is removed. The colony is thresholded to obtain the center point; if multiple possibilities are found, the user is asked to select the correct one. For ease of analysis and convolution, the program then carries out Cartesian-to-polar coordinate transformation via the MATLAB *scatteredInterpolant* object; radial features thus become horizontal in the 1000×1000 output images **(Fig. 1a-b)**. This “flattening” process takes advantage of the near-radial symmetry of the colonies. This custom script has enabled us to efficiently preprocess 1,000+ images thus far, and is easily used by students with little to no programming experience.

Two datasets were chosen from these images and further preprocessed separately **(Table 1)**. For colony segmentation, we selected a 306-image subset with a mix of colony area and background agar to aid training of deep learning models. Adaptive histogram equalization increased contrast, and any leftover Petri rim edge pixels were identified by thresholding and removed **(Fig. 1b5i)**. For ring boundary segmentation, we began with 558 images with distinct ring edges.

Images underwent adaptive histogram equalization, then smoothing with 13×13 Gaussian (standard deviation = 2) and 10×10 median filters **(Fig. 1b5ii)**.

### 2.2. Segmentation of swarm colony

We sought to develop a model which could output a colony mask given a preprocessed *P. mirabilis* image. We first developed an image processing-based algorithm for generating ground truths **(Fig. 1c)**. Various filters are applied to the preprocessed colony images and outputs are added, emphasizing faint edges; the result is thresholded to create a binary mask. In parallel, Gabor texture analysis generates a second option. If the areas of the two masks are similar, the first is used; otherwise, the user is asked to select the more complete mask. Next, morphological operations fill holes and eliminate small artifacts, creating an output ground truth mask. We removed 13 suboptimal mask-image pairs, leaving 306 pairs to split into training, validation, and test sets.

From each set, 128×128 patches were generated with stride 25 and labeled using the ground truths; the threshold for a positive patch was 50% foreground pixels. An image’s patch generation began at the top left and was stopped when white-space was reached, leading to about 212,000 training patches and 28,000 each for validation and testing. With total positive patches outweighing negative patches, class reweighting was used during training. The dataset was standardized by global mean subtraction.

Next, a CNN was trained to classify each given patch. We increased complexity from a single convolutional block until a final architecture with two convolutional layers was obtained: a convolutional layer with leaky ReLU activation followed by max pooling, a second such block with ReLU activation, flattening with Dropout of 0.2 immediately before and after a fully connected dense layer with ReLU activation, and a final classification dense layer with sigmoid activation. We also included augmentations of rotations, flips, and brightness changes, but observed that the model performed best without augmentation. Model hyperparameters included binary cross-entropy loss, Adam optimizer (learning rate 5e-4), and batch size 16. With early stopping, our model achieved 95% training and validation accuracy after 9 epochs.

To generate the colony segmentation, images were padded using reflection, then split into overlapping patches generated with stride 12; each patch was then classified using the trained CNN. Predictions were thresholded at 0.5. For each region in the original image, all patches overlapping it were identified. Predicted labels were then stacked on that region, creating a multichannel image in which each channel represents a specific overlapping prediction. Majority voting, i.e. labeling regions as positive if more than half of the overlapping predictions were positive, fused labels and generated the colony segmentation.

### 2.3. Segmentation of swarm ring boundaries

In parallel, we sought to segment ring boundaries (edges delineating periodic swarming phases) within *P. mirabilis* colonies. We developed another algorithm for generating ring ground truth masks on the preprocessed dataset. The Canny edge detector, using the derivative of a Gaussian filter with a 1.9 standard deviation and edge thresholds of 0.06 and 0.15, generates initial ring boundaries [18]. Retained edges are postprocessed with morphological-based methods, such as dilation, hit-miss operations, and skeletonization, resulting in binary output masks.

Whereas the previous CNN’s input was local patches, here a U-Net architecture was employed to consider the sparse yet localized and globally-dependent ring boundary pixels within a full-size image [14]. The ring boundary intricacies, which complicate annotation procedures, motivated an iterative supervised learning method **(Fig. 1d)**. In the first cycle, 558 preprocessed images and their Canny-derived masks were used for training, validation, and testing (with an 80-10-10% split) of a U-Net with a VGG-11 encoder pretrained on ImageNet [15, 16, 19, 20]. Hyperparameters included Dice loss, Adam optimizer (learning rate 1e-4), sigmoid activation, and batch size 3. Input images were padded to 1024×1024 using the border reflect method. After just 4 epochs, the model’s predictions on the unseen test set proved sufficient. Thus, this model was used to generate predictions on a broader collection of ∼750 images. Pixel probabilities were thresholded at 0.5 to yield binary masks.

For the second cycle, 300 predicted masks were skeletonized, then manually refined in ImageJ to connect broken boundaries and eliminate noise. The refined set was used to retrain the U-Net. Under early stopping (patience 3), an optimal model was obtained after 35 epochs when validation loss reached 0.23.

## 3. FINDINGS AND TESTING

In the process of arriving at our final pipelines, we evaluated various model architectures, hyperparameters, and postprocessing methods. For the colony segmentation task, only two convolutional layers and no augmentations were needed to successfully predict patch labels. Three label fusion methods were explored: majority voting, averaging of predictions, and a single convolutional layer with leaky ReLU activation. Although the single convolutional layer’s predicted masks were mostly accurate visually, their similarity to ground truth was less than those of the non-convolutional methods, suggesting a simpler method was better for fusing the predictions and that location of a given region within an image was not increasing labeling accuracy **(Fig. 2)**. The mean method was largely comparable to the majority voting method, but qualitatively the majority voting method appeared to generate the most accurate predictions at the colony edge, achieving a Dice score of 0.93 **(Table 2)**. In the important case of a barely visible outer ring of actively swarming bacteria, which is not fully captured by image processing and U-Net approaches, the patch-based majority voting approach successfully predicted the region as colony **(Fig. 2b, first row)**.

**Fig. 2.**
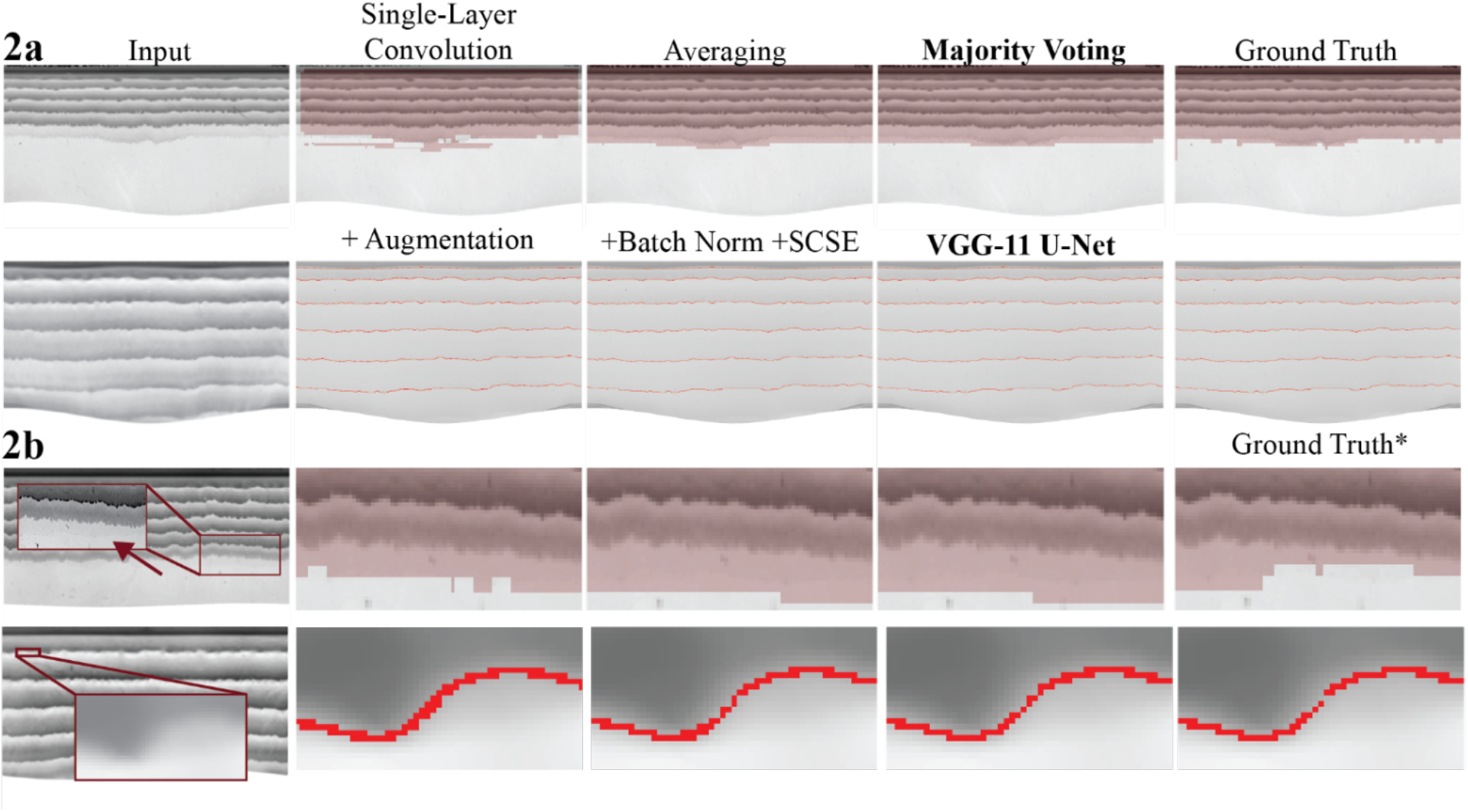
Qualitative segmentation predictions by various models on distinct test images. Predictions are compared to examples of **a**. high quality and **b**. suboptimal ground truths*. For the latter, cropped masks are enlarged to visualize where the pipeline exhibited better performance, by detecting a faint outer swarm front (red arrow) and connecting a broken ring boundary.

**Table 2.**
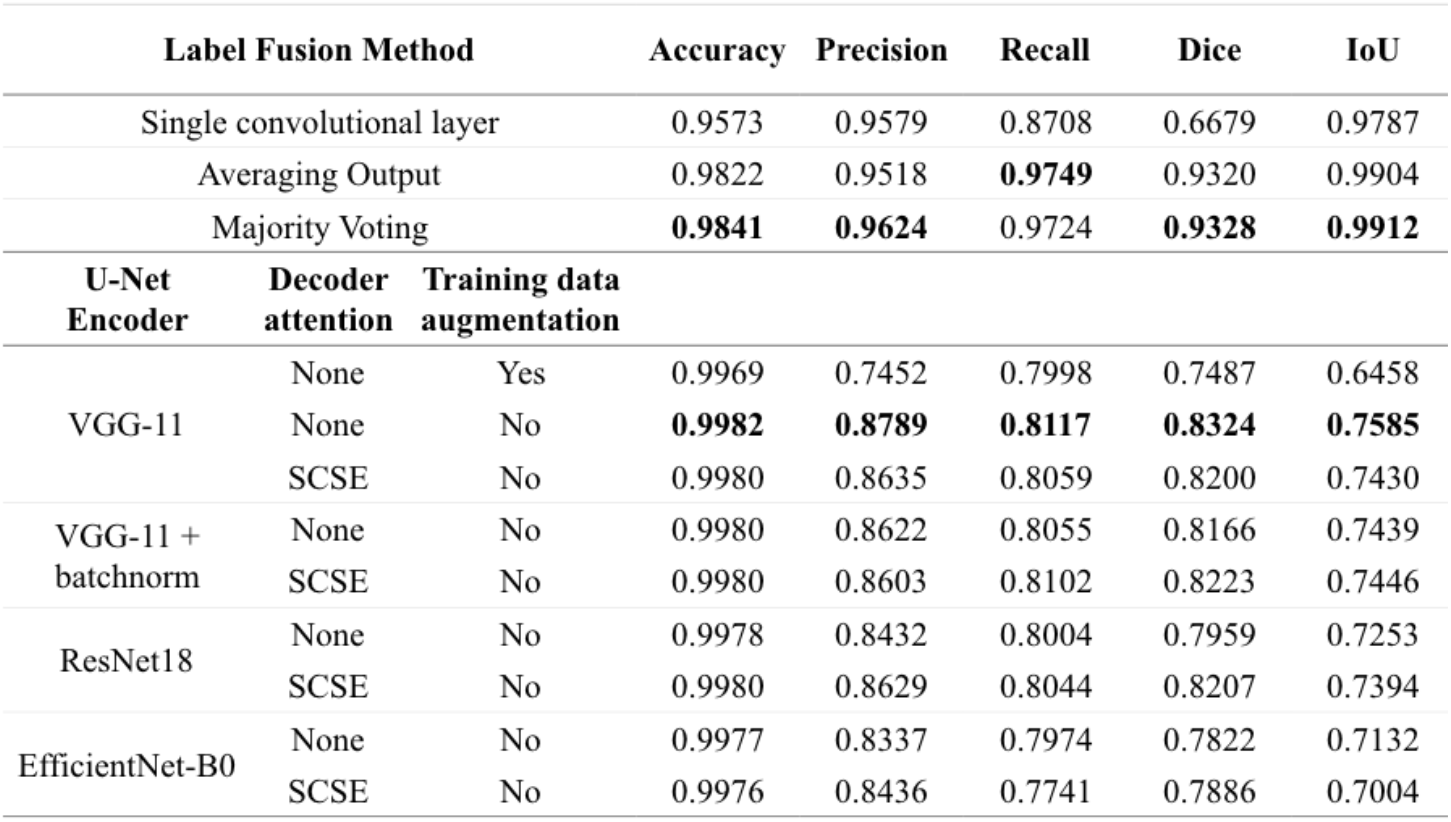
Performance of various approaches for segmentation.

While deep learning requires large datasets, biological data such as ours is laborious to generate and annotate. An important consideration was the number of initial predicted ring boundary masks to manually refine before the second training cycle. Various subsets of the manually refined masks (ranging from 8 to 300 images) were used to retrain the VGG-11 U-Net, with and without training data augmentations including rotations, flips, translations, and scaling. Decrease in validation loss became relatively marginal after 200 images, suggesting a dataset of 300 images was reasonable for this model. Training data augmentation resulted in overly thick predicted ring boundaries. Augmentations may have further amplified the dataset’s inherent underlying biological noise, impeding the model’s ability to precisely detect fine edges.

The un-augmented refined set was used to train the following additional encoders pretrained on ImageNet, with and without SCSE decoder attention: EfficientNet-B0, ResNet18, and VGG-11 with batch normalization [15, 19-23]. The addition of batch normalization and attention resulted in predictions qualitatively similar to those of the baseline VGG-11 U-Net **(Fig. 2)**. Ultimately, the baseline U-Net with a pretrained VGG-11 encoder yielded the best values for all test metrics, such as the highest Dice score of 0.83 and IoU of 0.76 **(Table 2)**. The model even improved upon certain ground truths. For example, the model connected a ring boundary that was erroneously disconnected by a user during the manual refinement step **(Fig. 2b, second row)**. Taken together, these results suggest that data augmentation and supplemental blocks within a network’s layers do not yield superior predictions for our task that would justify the computationally intensive additions.

Finally, we present a biologically and clinically relevant experiment to demonstrate the generalizability and utility of our pipeline **(Fig. 3)**. *P. mirabilis* colonies were grown at two standard laboratory conditions known to generate two colony patterns. Denser colonies with regular rings covering the whole plate grew on 1.5% agar in a 37**°**C incubator (Condition 1), while less dense colonies with a single ring covering part of the plate grew on 1.3% agar on the ∼25**°**C benchtop (Condition 2). The pipeline was used to predict colony and ring boundary masks for each image. Predicted colony masks were used to calculate colony area. As the VGG-11 U-Net occasionally missed the faintest regions of boundaries, predicted ring boundary masks were postprocessed to omit image columns which lacked the true number of ring boundaries. Maximum inter-ring-boundary distances were measured. A paired t-test demonstrated significant difference of these colony features between the two conditions, with p<<0.05 for both. These features, distinguishing the conditions, and others could be later used to evaluate motility of clinical or experimentally relevant strains. This experiment demonstrates how our pipeline can be adapted to distinguish different colony patterns that researchers might encounter when working with swarming bacteria. Quantitative findings from experiments such as this have clinical significance, as surface hardness, environmental temperature, and nutrient availability affect swarming motility, and thus are of interest to understand pathogenicity.

**Fig. 3.**
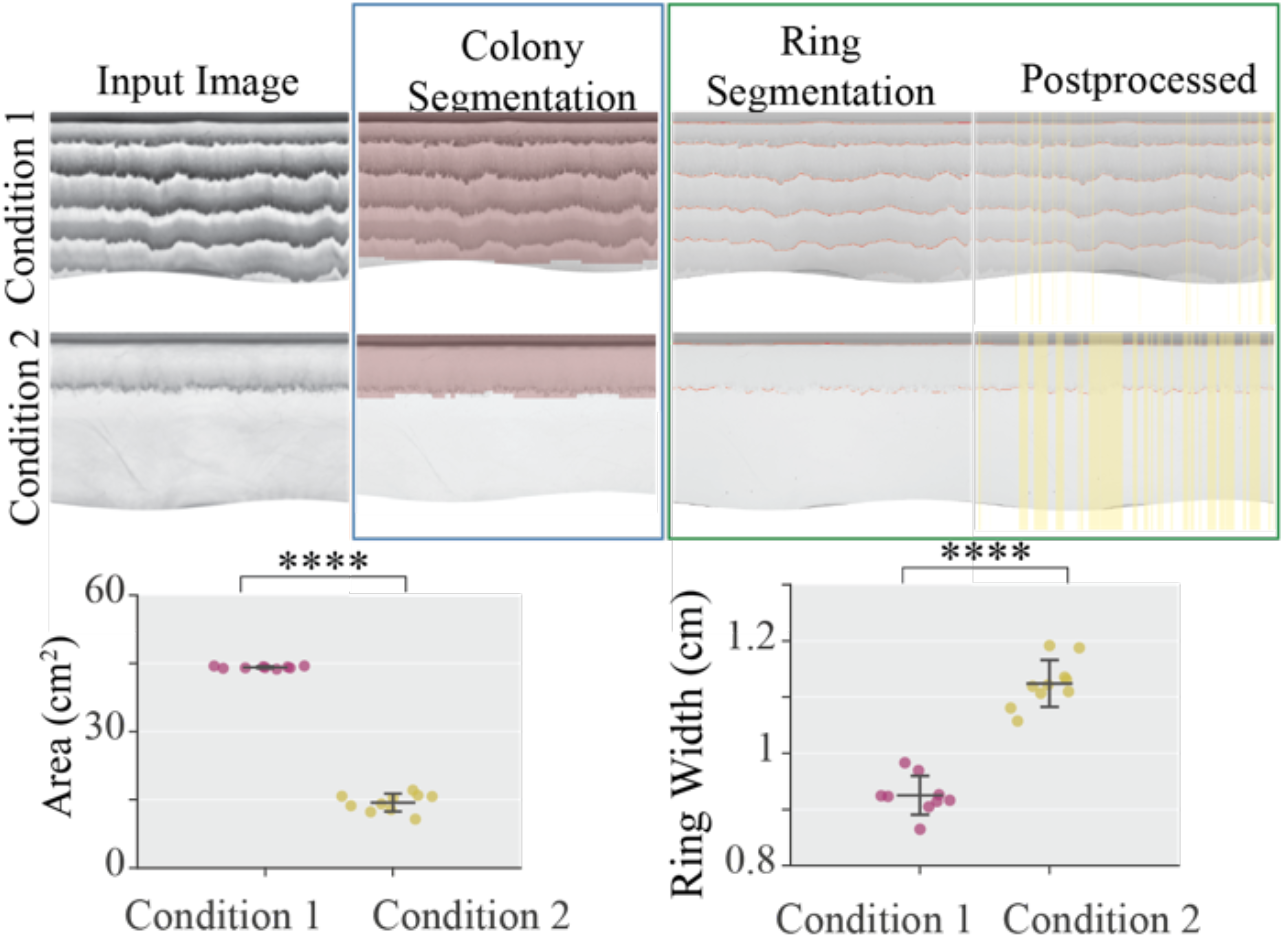
Example application of pipeline. Representative images are shown of bacteria grown on 1.5% agar at 37°C (9 images, Condition 1) and 1.3% agar at 25°C (10 images, Condition 2). Ring boundary masks were postprocessed to retain only relevant information prior to feature extraction. Error bars are STD. For colony area, mean at Condition 1 was 44.1 cm^2^ and STD = 0.23; mean at Condition 2 was 14.4 cm^2^ and STD = 1.98. p = 4.53e-19. For ring width, the maximum distance between any two ring boundaries on the plate was calculated; for Condition 1 the mean was 0.95 cm with STD = 0.03 and for Condition 2 the mean was 1.12 cm, STD = 0.04. p = 2.72e-9.

## 4. CONCLUSION

We have developed a dual-task pipeline for accurately segmenting motility-dependent macroscopic colonies and ring boundaries within images from *P. mirabilis* swarming assays. Colony segmentation captures faint active swarm rings and enables evaluation of overall colony features. Ring boundary segmentation allows quantification of colonies’ repetitive pattern features, which have thus far not been analyzed in detail. Easing the burden of manual input, our pipeline includes preprocessing, data compilation, postprocessing, and feature extraction functions which are easily scaled to thousands of images, and can enable researchers to collect and analyze larger datasets. At the same time, our patch-based and transfer learning approaches allowed us to work with biological datasets that are small relative to typical deep learning datasets. Overall, the pipeline provides essential information to analyze *P. mirabilis* motility. In the future, it could be applied to analyze the motility and macroscopic colonies of other clinical isolates and soil microbes with more complex features such as branched and fractal structures. We plan to integrate this pipeline into a single package such as an ImageJ plugin for dissemination. This work can serve as a framework for researchers developing new computational tools to analyze bacteria with diverse colony morphologies and roles in infectious disease spreading.

## 5. COMPLIANCE WITH ETHICAL STANDARDS

This is a computational study for which no ethical approval was required.

## 6. ACKNOWLEDGMENTS

This work was supported by an NSF CAREER Award (1847536), Blavatnik Fund for Innovations in Health (T.D.), and National Science Foundation Graduate Research Fellowship (A.D.). *Proteus mirabilis* (ATCC 7002) was kindly provided by Dr. Philip Rather.

## 8. DATA AND CODE AVAILABILITY

All code not in the Github and all data are proprietary and managed by the Columbia Technology Ventures Office of Intellectual Property. They are available from the corresponding author T.D. upon reasonable request, after permission from the Columbia Technology Ventures Office of Intellectual Property.

